# *Neotrygon indica* sp. nov., the Indian-Ocean blue spotted maskray (Myliobatoidei, Dasyatidae)

**DOI:** 10.1101/179911

**Authors:** Annam Pavan-Kumar, Rajan Kumar, Pranali Pitale, Kang-Ning Shen, Philippe Borsa

**Affiliations:** Fish Genetics and Biotechnology Division, ICAR-Central Institute of Fisheries Education, Mumbai — 61, India; Fisheries Resources and Postharvest Division, ICAR-Central Institute of Fisheries Education, Mumbai, India; Aquatic Technology Laboratories, Agricultural Technology Research Institute, Taiwan, Republic of China; Institut de recherche pour le développement (IRD), UMR 250 “Ecologie marine tropicale des océans Pacifique et Indien”, Nouméa, New Caledonia

## Abstract

The blue-spotted maskray, previously N *kuhlii*, consists of up to eleven lineages representing separate species. Nine of these species (N *australiae, N. bobwardi, N. caeruleopunctata, N. malaccensis*, N *moluccensis*, N *orientale, N. vali*, N *varidens*, N *westpapuensis)* have already been formally described and two (Indian-Ocean maskray and Ryukyu maskray) remain undescribed. Here the Indian-Ocean maskray is described as a new species, *Neotrygon indica* sp. nov.

---

To describe species, taxonomists typically use morphology as primary evidence and increasingly often, genetic information to support morphology ^1^. Although genetics have proven essential to delineate species in many instances, genetic information such as allele frequencies or DNA sequences is still exceptionally used as primary evidence in new species descriptions, in part if not mostly because of considerable peer pressure against DNA-based descriptions ^1^. However, there is no objective reason to dismiss genetics as the primary source of characters for species delineation, description, and identification ^1-3^.

The blue-spotted maskray, previously *N. kuhlii* (Müller and Henle 1841), has a wide Indo-West Pacific distribution, from eastern Africa to Japan ^4-7^. It was hypothesized to be a species complex after a barcoding survey revealed several deeply divergent mitochondrial lineages ^8^. The hypothesis of a species complex was subsequently validated ^6,9^. This species complex consists of up to eleven parapatrically-distributed lineages representing separate species ^6,9-12^ of which nine have already been formally described ^7,12-14^. These are *N. australiae* Last, White and Séret 2016, *N. bobwardi* Borsa, Arlyza, Hoareau and Shen 2017, *N. caeruleopunctata* Last, White and Séret 2016, *N. malaccensis* Borsa, Arlyza, Hoareau and Shen 2017, *N. moluccensis* Borsa, Arlyza, Hoareau and Shen 2017, *N. ońentale* Last, White and Séret 2016, *N. vali* Borsa 2017, *N. varidens* (Garman 1885), and *N. westpapuensis* Borsa, Arlyza, Hoareau and Shen 2017. Two other species in the genus, the nominal *N. kuhlii* from Vanikoro and *N. trigonoides* possess distinctive spot patterns ^3,15,16^ that tell them apart from the blue-spotted maskray as it was originally described by J. Müller and F.G.J. Henle ^4^. Based on the only available information on colour patterns, one cannot exclude that *N. kuhlii* as it has been redefined recently ^14^ and *N. trigonoides* are synonyms ^16^.

Some authors have attempted to use morphological characters as primary information for the description of cryptic species in the blue-spotted maskray complex ^14^ and a recent revision has ostensibly ignored genetic evidence even though not a single morphological character among those utilized for the description or redescription of four species in the complex ^14^ was undisputably diagnostic of any of the species ^7^. Also, the apportion of environmental vs. genetic determination in these morphological characters had not been evaluated ^7^. Morphological diagnoses provided so far for the blue-spotted maskray ^14^ are therefore hardly relevant, as previously mentioned for the *Himantura uarnak* complex, another species complex among stingrays ^3^. In contrast, DNA sequences offer a profusion of diagnostic characters in stingrays ^3,6,9,14,18^. Species-specific haplogroups were observed in the blue-spotted maskray, based on DNA sequences ^3,9^.

The objective of the present paper is to formally describe the Indian-Ocean maskray, primarily based on the mitochondrial DNA sequences of fresh specimens from the eastern coast of India and of previously-reported material from India and Tanzania.

## Materials and Methods

### Material examined

Eleven specimens of the new species that were examined for spot patterns are listed in Table 1. Specimen NKGMF-3 from the Gulf of Mannar, January 2017, to be designated as the holotype of the new species, was deposited at the Marine Biology Regional Centre (MBRC) in Chennai, India. The MBRC is part of the network of museums managed by, and an official repository of the Zoological Survey of India. Specimen NKGMF-1 from the Gulf of Mannar, January 2017, to be chosen as paratype of the new species, was deposited at the Fish Genetics and Biotechnology (FGB) laboratory, ICAR-Central Institute of Fisheries Education (CIFE) in Mumbai, India. Voucher material of the new species also includes a specimen collected by APK and colleagues in Visakhapatnam, Andhra Pradesh, Bay of Bengal on 15 August 2011. The specimen was discarded but a sub-sample of tissue was registered at the Central Institute of Fisheries Education, Mumbai under no. VIZNK-01. A photograph of this individual is available from the Barcoding of Life Data systems database (BOLD; http://www.barcodinglife.com/) ^20^ under accession no. BOLD:ACB9305. Photographs of blue-spotted maskrays from India deposited in FishBase ^21^ were not retained for the analysis of spot patterns because of insufficient resolution.

**Table 1.** Spot patterns on the dorsal side of 41 blue-spotted maskray (*Neotrygon* spp.) specimens sorted by species. *L*, left pectoral fin; *R*, right pectoral fin. *Occipital mark*, diffuse dark blotch at rear end of neurocranium that was categorized as either absent (*0*), weak (*1*) or conspicuous (*2*).

| Species,<br>Specimen no. (field no.) | Locality of collection,<br>country | N ocellated blue spots |  |  | N dark<br>speckles<br>(<1% DW)<br>L, R | N dark<br>spots<br>(>1% DW)<br>L, R | Occipital<br>mark |
| --- | --- | --- | --- | --- | --- | --- | --- |
|  |  | 1. Small<br>L, R | 2. Medium<br>L, R | 3. Large<br>L, R |  |  |  |
| *N. australiae* |  |  |  |  |  |  |  |
| CSIRO 7016-01 | Weipa, Australia | 14, 12 | 17, 20 | 3, 0 | 8, 9 | 0, 0 | 0 |
| MZB-20863 | Tanjung Sulamo, Indonesia | 25, 25 | 28, 32 | 1, 0 | 13, 6 | 0, 0 | 0 |
| *N. bobwardi* |  |  |  |  |  |  |  |
| MZB-20843 | Meulaboh, Indonesia | 28, 29 | 13, 11 | 0, 0 | 16, 16 | 0, 0 | 0 |
| 20150524-D | Padang, Indonesia | 5, 11 | 4, 2 | 0, 0 | 6, 6 | 0, 0 | 0 |
| 20150524-E | Padang, Indonesia | 16, 11 | 9, 8 | 0, 0 | 8, 15 | 0, 0 | 2 |
| 20150524-F | Padang, Indonesia | 20, 36 | 14, 8 | 0, 0 | 3, 0 | 0, 0 | 1 |
| 20150524-11 | Padang, Indonesia | 8, 8 | 2, 1 | 0, 0 | 6, 13 | 0, 0 | 2 |
| *N. caeruleopunctata* |  |  |  |  |  |  |  |
| CSIRO H 7850-01<sup>b</sup> | Sadeng, Indonesia | 25, 22 | 4, 1 | 0, 0 | 17, 30 | 0, 0 | 1 |
| 20080131_BL (NK-K-BL) | Bali, Indonesia | 21, 24 | 6, 5 | 0, 0 | 43, 54 | 0, 0 | 1 |
| MZB-22131 (BAL-S) | Bali, Indonesia | 31, 22 | 5, 9 | 0, 1 | 0, 1 | 0, 0 | 1 |
| *N. indica* sp. nov. |  |  |  |  |  |  |  |
| ZSI/MBRC/F.1495 | Gulf of Mannar, India | 9, 10 | 0, 0 | 0, 0 | 19, 34 | 3, 2 | 2 |
| CIFEFGB/NKGM1 | Gulf of Mannar, India | 27, 26 | 0, 0 | 0, 0 | 37, 22 | 4, 1 | 2 |
| VIZNK-01 | Visakhapatnam, India | 3, 12 | 1, 9 | 0, 0 | 13, 10 | 0, 0 | 2 |
| NKTTK-1 | Tuticorin, India | 30, 27 | 3, 3 | 0, 0 | 90, 86 | 0, 1 | 2 |
| NKTTK-2 | Tuticorin, India | 20, 8 | 0, 0 | 0, 0 | 24, 32 | 0, 1 | 2 |
| NKGMM-2 | Gulf of Mannar, India | 24, 26 | 7, 4 | 0, 0 | 30, 33 | 4, 1 | 2 |
| NKGMM-4 | Gulf of Mannar, India | 14, 18 | 8, 4 | 0, 0 | 40, 38 | 5, 6 | 2 |
| NITT-1 | Tuticorin, India | 28, 28 | 20, 14 | 0, 0 | 68, 73 | 1, 1 | 2 |
| NITT-3 | Tuticorin, India | 10, 32 | 9, 3 | 0, 0 | 34, 35 | 1, 3 | 2 |
| ZAN 1 | Zanzibar, Tanzania | 44, 51 | 29, 21 | 2, 2 | 10, 15 | 0, 0 | 0 |
| IRD-20170720-A | Zanzibar, Tanzania | 32, 35 | 6, 6 | 0, 0 | 16, 21 | 1, 0 | 2 |
| *N. malaccensis* |  |  |  |  |  |  |  |
| MZB-20847 | Kuala Lama, Indonesia | 2, 4 | 14, 10 | 0, 1 | 5, 3 | 0, 0 | 0 |
| *N. moluccensis* |  |  |  |  |  |  |  |
| MZB-20866 | Tual, Indonesia | 36, 30 | 9, 9 | 0, 0 | 53, 53 | 0, 1 | 0 |
| MZB-20864 | Ambon, Indonesia | 9, 2 | 7, 12 | 0, 0 | 5, 2 | 0, 0 | 1 |
| *N. orientale* |  |  |  |  |  |  |  |
| CSIRO H7858-01<sup>a</sup> | Muara Kintap, Indonesia | 20, 14 | 15, 17 | 1, 1 | 2, 1 | 0, 0 | 0 |
| MZB-20852 | P. Pabelokan, Indonesia | 5, 9 | 8, 3 | 1, 0 | 21, 17 | 0, 0 | 0 |
| MZB-20851 | Pulau Pari, Indonesia | 22, 18 | 9, 6 | 2, 2 | 12, 8 | 0, 0 | 0 |
| MZB-20850 | Pulau Peniki, Indonesia | 12, 22 | 12, 11 | 2, 0 | 2, 1 | 0, 0 | 1 |
| BO424 | Tanjung Manis, Malaysia | 35, 41 | 22, 6 | 0, 0 | 16, 13 | 0, 0 | 0 |
| WJC-5367 | Kota Kinabalu, Malaysia | 17, 19 | 10, 8 | 0, 1 | 8, 4 | 0, 0 | 0 |
| WJC-5368 | Kota Kinabalu, Malaysia | 19, 20 | 6, 12 | 0, 0 | 6, 5 | 0, 0 | 0 |
| WJC-5369 | Kota Kinabalu, Malaysia | 20, 23 | 14, 10 | 0, 0 | 3, 2 | 0, 0 | 0 |
| *N. vali* |  |  |  |  |  |  |  |
| CSIRO H7723-01 | Honiara, Solomon Is. | 2, 4 | 1, 1 | 0, 0 | 3, 6 | 0, 0 | 0 |
| Randall<sup>19</sup> | Solomon Is. | 11 | 4 | 0 | 6 | 0 | 0 |
| Rosenstein<sup>12</sup> | Mbike wreck, Solomon Is. | 21 | 6 | 0 | 7 | 0 | 0 |
| *N. varidens* |  |  |  |  |  |  |  |
| ASIZP0073604 | Taiwan | 0, 1 | 0, 0 | 0, 0 | 0, 0 | 0, 0 | 1 |
| ASIZP0073605 | Taiwan | 3, 2 | 0, 0 | 0, 0 | 0, 0 | 0, 0 | 1 |
| ASIZP0073535 | Taiwan | 3, 1 | 0, 0 | 0, 0 | 0, 0 | 0, 0 | 1 |
| ASIZP0078458 | Taiwan | 7, 16 | 0, 0 | 0, 0 | 0, 0 | 0, 0 | 1 |
| ASIZP0805866 | Taiwan | 4, 3 | 5, 6 | 0, 0 | 2, 7 | 0, 0 | 1 |
| *N. westpapuensis* |  |  |  |  |  |  |  |
| MZB-20867 | Biak, West Papua | 41, 37 | 25, 13 | 0, 2 | 10, 7 | 0, 0 | 1 |

Five specimens, including four from India and one from Tanzania were measured ^14^ using a ribbon meter, a ruler, and a vernier caliper. Three of these, including the holotype and the paratype of the new species were X-ray photographed. Specimens of the new species that were characterized by their nucleotide sequences at loci *CO1* and *cytochrome b* are listed in Table 2 and include 13 specimens from India and 6 specimens from Tanzania. Comparative material for the analysis of nucleotide sequences included blue-spotted maskray specimens listed previously ^9^ and an additional specimen from Taiwan deposited in the fish collections of Academia Sinica, Taipei, and registered under no. ASIZP0805866.

**Table 2.** *Neotrygon indica* sp. nov. specimens examined for nucleotide variation at mitochondrial loci *CO1* and *cytochrome b*.

| Specimen no. (field no.) | Locality of collection, country | GenBank accession no. |  | Source |
| --- | --- | --- | --- | --- |
|  |  | *CO1* | *cytochrome b* |  |
| NBFGR:CHN 156 | Chennai, India | HM467799 | - | Mined from GenBank |
| NBFGR:CHN:R24 | Chennai, India | KF899609 | - | Mined from GenBank |
| NBFGR:CHN:R25 | Chennai, India | KF899610 | - | Mined from GenBank |
| NBFGR:CHN:R165 | Chennai, India | KF899611 | - | Mined from GenBank |
| NBFGR:CHN:NK13 | Chennai, India | KF899612 | - | Mined from GenBank |
| NBFGR:CHN:199 | Chennai, India | KF899613 | - | Mined from GenBank |
| - | Tamil Nadu, India | KR003770 | - | Mined from GenBank |
| ZSI/MBRC/F.1495 | Gulf of Mannar, India | MG017600 | MG017605 | Present study |
| NKGMM-2 | Gulf of Mannar, India | MG017601 | MG017606 | Present study |
| NKVSKP-1 | Visakhapatnam, India | JX978329, | MG017607 | Mined from BOLD, |
|  |  | MG017602 |  | Present study |
| NKVSKP2 | Visakhapatnam, India | MG017603 | MG017608 | Present study |
| NKVSKP3 | Visakhapatnam, India | MG017604 | MG017609 | Present study |
| NKTTK1 | Tuticorin, India | - | MG017610 | Present study |
| ZANZ 1 | Zanzibar, Tanzania | JX263421 | KU497907 | Ref. 10 |
| 80611 | Tanga, Tanzania | KC249906 | - | Ref. 6 |
| KNS-TZN1-52 | Pemba Island, Tanzania | KU498035 | KU497908 | Ref. 9 |
| KNS-ZAN3-86 | Pemba Island | KU498036 | KU497909 | Ref. 9 |
| KNS-ZAN4-71 | Pemba Island | KU498037 | KU497910 | Ref. 9 |
| KNS-ZAN5-80 | Pemba Island | KU498038 | KU497911 | Ref. 9 |

### Spot pattern analysis

The diameter of ocellated blue spots on the dorsal side of the left and right pectoral fins, relative to disc width (DW), was measured from photographs ^7,12,14,15^; Figs. 1 and 2). Ocellated blue spots were qualified as “small” when their maximum diameter was ≤ 2% DW, “medium” when ≤ 4% DW and “large” when > 4% DW ^15^. Dark speckles (≤ 1% DW) and dark spots (> 1% DW) were also counted on the dorsal surface of the disc. The counts did not include those speckles and spots located within the dark band around eyes that forms the mask ^15^. The presence or absence of a darker occipital mark at the rear end of the neurocranium was also checked and categorized as “absent”, “weak”, or “conspicuous”. Spot patterns were compared among individuals through correspondence analysis (CA) ^22^. CA was run using the FacTOMineR package ^23^ under R ^24^.

**Fig. 1.**
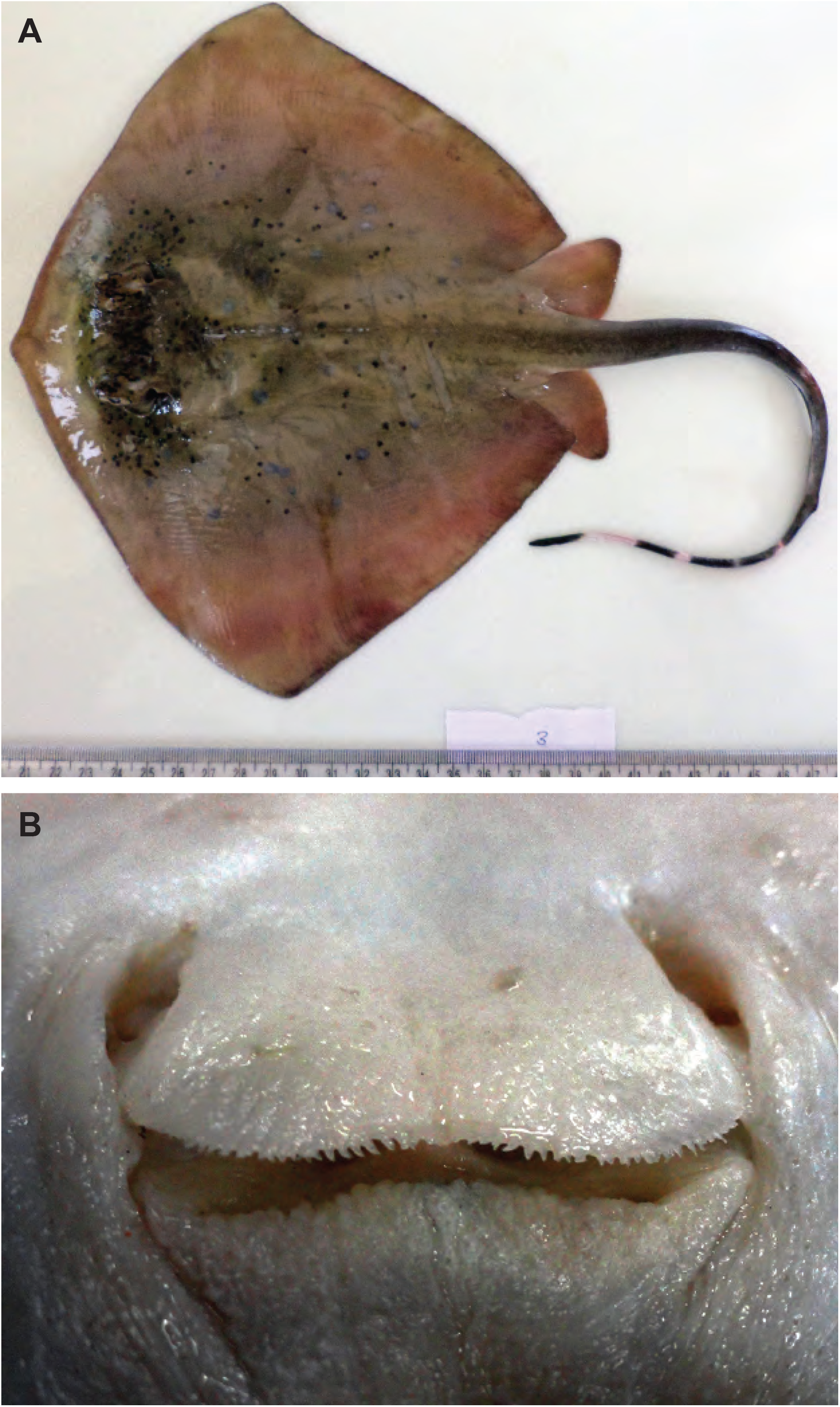
Holotype of *Neotrygon indica* sp. nov.: female specimen, 174 mm disc length from the Gulf of Mannar, Tamil Nadu, India (09.12°N 79.46°E) registered under no. ZSI/MBRC/F.1495 at the Marine Biology Regional Centre in Chennai, India. **A.** Dorsal side (photograph by RK). **B.** Oronasal region (photograph by APK).

**Fig. 2.**
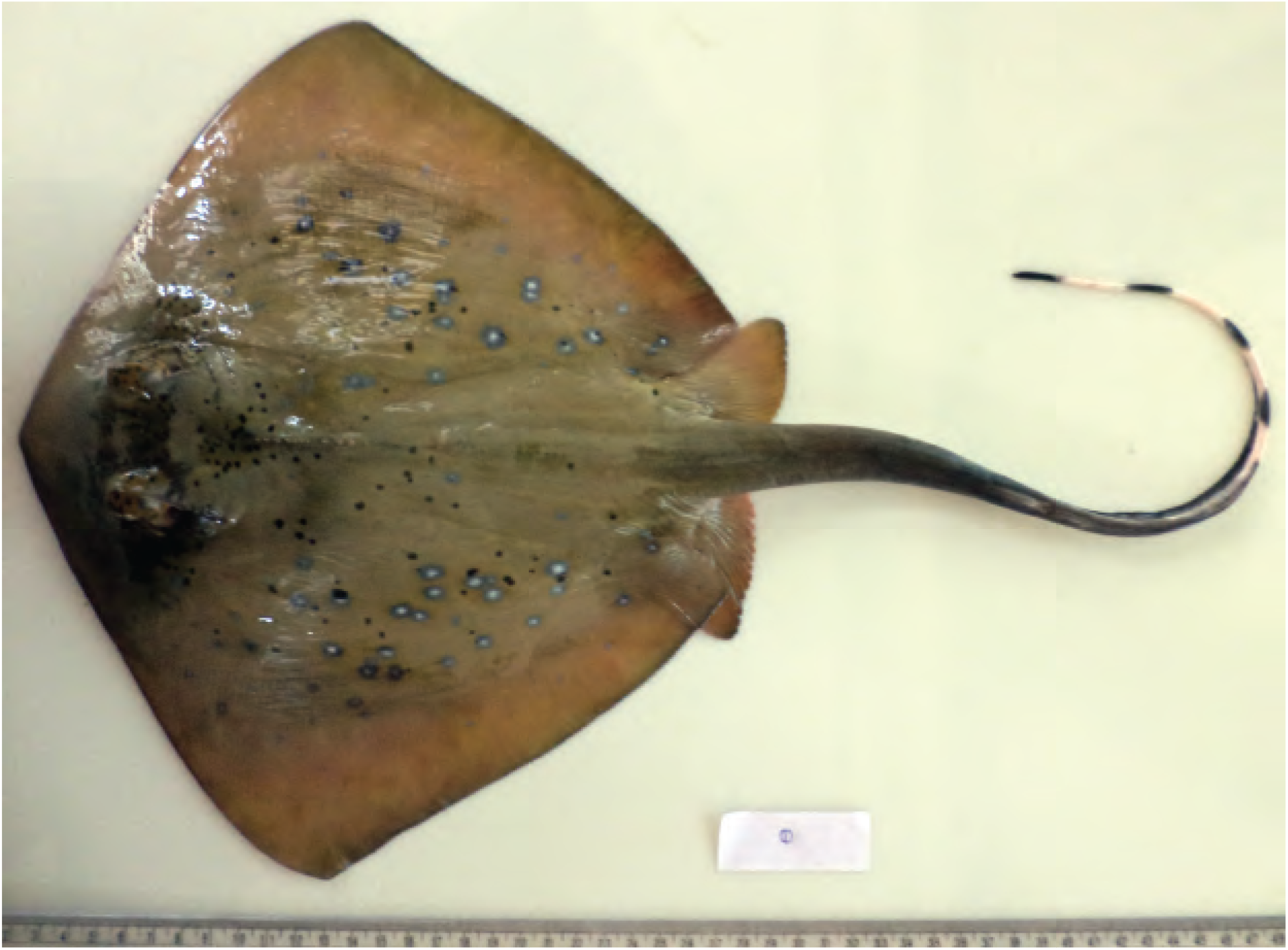
Paratype of *Neotrygon indica* sp. nov.: female specimen no. CIFEFGB/NKGM1, 182 mm disc length from the Gulf of Mannar, Tamil Nadu, India (9.12°N 79.46°E) (photograph by RK).

### Analysis of nucleotide sequences

The DNA of six specimens of the new species and of specimen ASIZP0805866 from Taiwan was extracted using the phenol-chloroform protocol and the DNEasy extraction kit (Qiagen GmbH, Hilden, Germany), respectively. A fragment of each the *CO1* and the *cytochrome b* genes was amplified by polymerase chain reaction according to previously published protocols ^9^. Nucleotide sequencing was done in both forward and reverse directions using the Sanger method. The consensus sequences were obtained after assembling the direct and reverse traces under BioEdiT v. 7.1.11 ^25^. Sequences were then edited and aligned under BioEdiT. We chose as reference for numerotating nucleotides the complete mitogenome sequence of *Neotrygon ońentale* ^26^, accessible from GenBank (http://www.ncbi.nlm.nih.gov) under no. KR019777. Five out of 11 *N. indica* sp. nov. individuals characterized by their spot patterns (Table 1) had their nucleotide sequences at loci *CO1* and/or *cytochrome b* examined (Table 2).

The phylogeny of concatenated *CO1* + *cytochrome b* gene haplotypes was inferred using the maximum-likelihood (ML) method under Mega6 ^27^. The most likely nucleotide-substitution model, which was determined according to the Bayesian information criterion, was the Tamura-Nei model (TN93) ^28^ where a discrete Gamma distribution (Γ = 0.76) was used to model evolutionary rate differences among nucleotide sites and invariable sites were allowed. The ML tree was rooted by choosing New Caledonian maskray *N. trigonoides* as outgroup ^15^. The robustness of nodes in the tree was tested by bootstrap resampling.

Based on the larger number of *CO1* gene sequences available (Table 2), a parsimony network was constructed to further explore the relationships among haplotypes of the new species, and with those of the geographically adjacent species *N. bobwardi, N. caeruleopunctata* and *N. malaccensis.* Median-joining parsimony analysis was done using NETWORK ^29^ on the nucleotide sequence matrix of 57 individual *CO1* gene haplotypes compiled from the present work and from previous reports ^9^. This matrix comprised sequences from blue-spotted maskrays from India (N = 11) and Tanzania (N = 5) as well as from the geographically adjacent *N. bobwardi* from the Andaman Sea and Western Sumatera (N = 13), *N. caeruleopunctata* from southern Java and southern Bali (N = 18), and *N. malaccensis* from the Malacca Strait and the eastern Andaman Sea (N = 10). Prior to median-joining analysis, the sequence matrix was trimmed to a core length of 638 bp, between nucleotide sites 68 and 705 of the *CO1* gene. The placement of the root was inferred from the maximum-likelihood tree of concatenated *CO1* + *cytochrome-b* gene sequences. Parsimony network analysis was similarly run on a matrix of 48 *cytochrome b* gene sequences, trimmed to a core length of 795 bp, between nucleotide sites 238 and 1032 of the *cytochrome b* gene. This comprised six sequences from India and five sequences from Tanzania (Table 2), as well as seven *N. bobwardi*, 17 *N. caeruleopunctata*, and 13 *N. malaccensis*.

Mean nucleotide distances within and between species were estimated from the same 57-individual *CO1* and 48-individual *cytochrome-b* sequence datasets using Mega6. According to the Bayesian information criterion, the best substitution model for the *CO1* gene sequence dataset was K2+G (*Γ* = 0.07). The best substitution model for the *cytochrome b* sequence data was HKY+G (*Γ* = 0.05), but the model used for the calculations was the second-best (TN93+G) which, unlike the former, is available in Mega6.

### Barcode index number assignment

The BOLD datasystems distinguishes clusters of sequences that qualify as operational taxonomic units, i.e. putative species using the Refined Single Linkage algorithm ^3σ^. The latter “clusters sequences with high similarity and connectivity and separates those with lower similarity and sparse connectivity”. Each putative species thus flagged is allocated a unique barcode index number (BIN) in BOLD. Borsa et al. ^7^ have established the homology of BIN numbers with cryptic blue-spotted maskray lineages by visual inspection of the placement of the *CO1* gene sequences retrieved from BOLD in a reference maximum-likelihood tree rooted by *N. trigonoides*.

### Notice

The present article in portable document (.pdf) format is a published work in the sense of the International Code of Zoological Nomenclature ^31^ or Code and hence the new names contained herein are effectively published under the Code. This published work and the nomenclatural acts it contains have been registered in ZooBank (http://zoobank.org/), the online registration system for the International Commission on Zoological Nomenclature. The ZooBank life science identifier (LSID) for this publication is urn:lsid:zoobank.org:pub:10551DF9-4B93-40ED-8910-012C5DCF8B96. The information associated with this LSID can be viewed through any standard Internet browser by appending the LSID to the prefix “http://zoobank.org/”. The online version of the present paper is archived and available from the *bioRxiv*, Cold Spring Harbor NY, U.S.A. (http://www.biorxiv.org/) and *haL-IRD*, France (http://www.hal.ird.fr/) repository websites.

## Results and Discussion

### Spot patterns

Spot patterns of 11 *N. indica* sp. nov. specimens were summarized and compared to nine other species of the blue-spotted maskray complex (Table 1). *N. indica* sp. nov. specimens from India were characterized by a moderately large number of small ocellated blue spots (*n* = 15-57), a generally low number of medium-sized ocellated blue spots (*n* = 0-34), a total absence of large ocellated blue spots, a high number of dark speckles (*n* = 23-176), generally a few dark spots (*n* = 0-11; only exceptionally present in the other species examined), and a conspicuous occipital mark. These features were visible on the holotype (Fig. 1) and the paratype (Fig. 2) of the new species. One of the two individuals from Tanzania had similar features, but not the other (Table 1). A CA run focused on the specimens from the Indian Ocean formed a cluster mostly distinct from the other species (Fig. 3). *Neotrygon indica* sp. nov. specimens tended to have a higher count of dark spots and dark speckles than the other three species (Fig. 3).

**Fig. 3.**
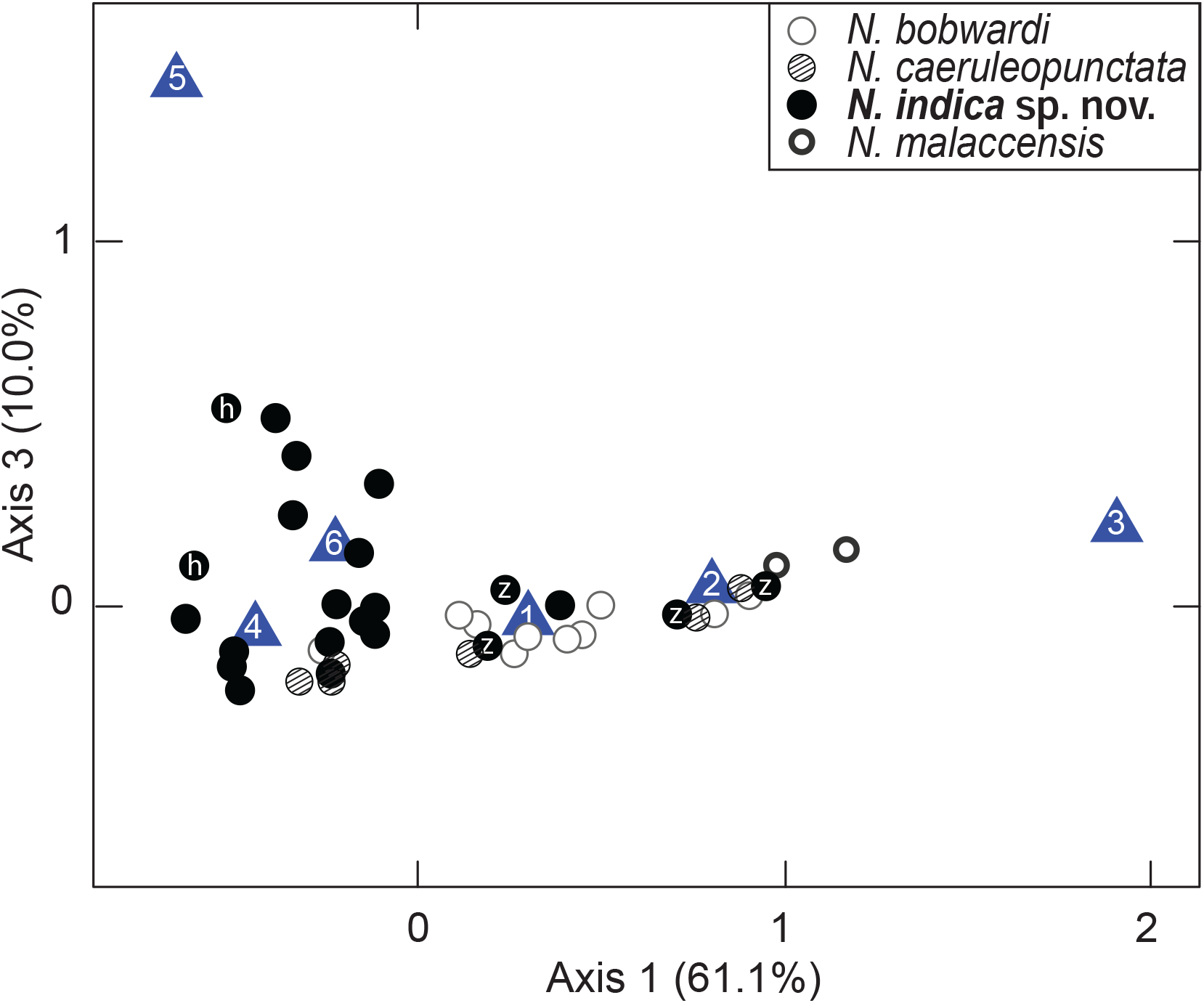
Correspondence analysis: projection of individuals characterized by counts of ocellated blue spots and dark spots and speckles, and by intensity of occipital mark on the dorsal side of a half-disc. *Blue triangles* indicate the position of the variables used to characterize spot patterns: *1*. Small ocellated blue spots; *2*. Medium-sized ocellated blue spots; 3. Large ocellated blue spots; *4*. Dark speckles; *5*. Dark spots; *6*. Occipital mark. *h* holotype; *z* sample from Zanzibar.

As shown by our results, some overlap in spot patterns was observed between some pairs of species. Also, one expects that with increased geographic sampling, based on the current sample sizes, more variation will show up within a species, hence further blurring the distinction between species and further relativizing diagnoses based on spot patterns.

### Genetic distinctness of the Indian-Ocean blue-spotted maskray

It has been recently reported that blue-spotted maskray populations from the Indian Ocean west of Bali (including *N. bobwardi* and *N. indica* sp. nov.) were “unresolved” but hypothesized that they were closely related to *N. caeruleopunctata* ^14^. Also, a single same BIN number (BOLD: AAA5611) has so far been allocated by BOLD to *N. caeruleopunctata, N. indica* sp. nov. and *N. malaccensis* ^7^ suggesting that the level of genetic differentiation at the *CO1* locus is not sufficiently clearcut to raise the three species, together with the closely related *N. bobwardi* to the level of fully distinct operational taxonomic units in the sense usually understood by the barcoding community. More in-depth investigation of the genetic relationships among populations within this group of geographically adjacent species from the Indian Ocean was warranted. Here, *Neotrygon indica* sp. nov. haplotypes formed a distinct haplogroup on the maximum-likelihood phylogenetic tree of concatenated *CO1*+*cytochrome b* gene sequences (Fig. 4), confirming previous results ^9^. However, it cannot be excluded that the Indian-Ocean maskray comprises at least three subclades, including two from India and one from Tanzania. Testing this hypothesis would require more comprehensive and denser sampling of the wide Indian Ocean.

**Fig. 4.**
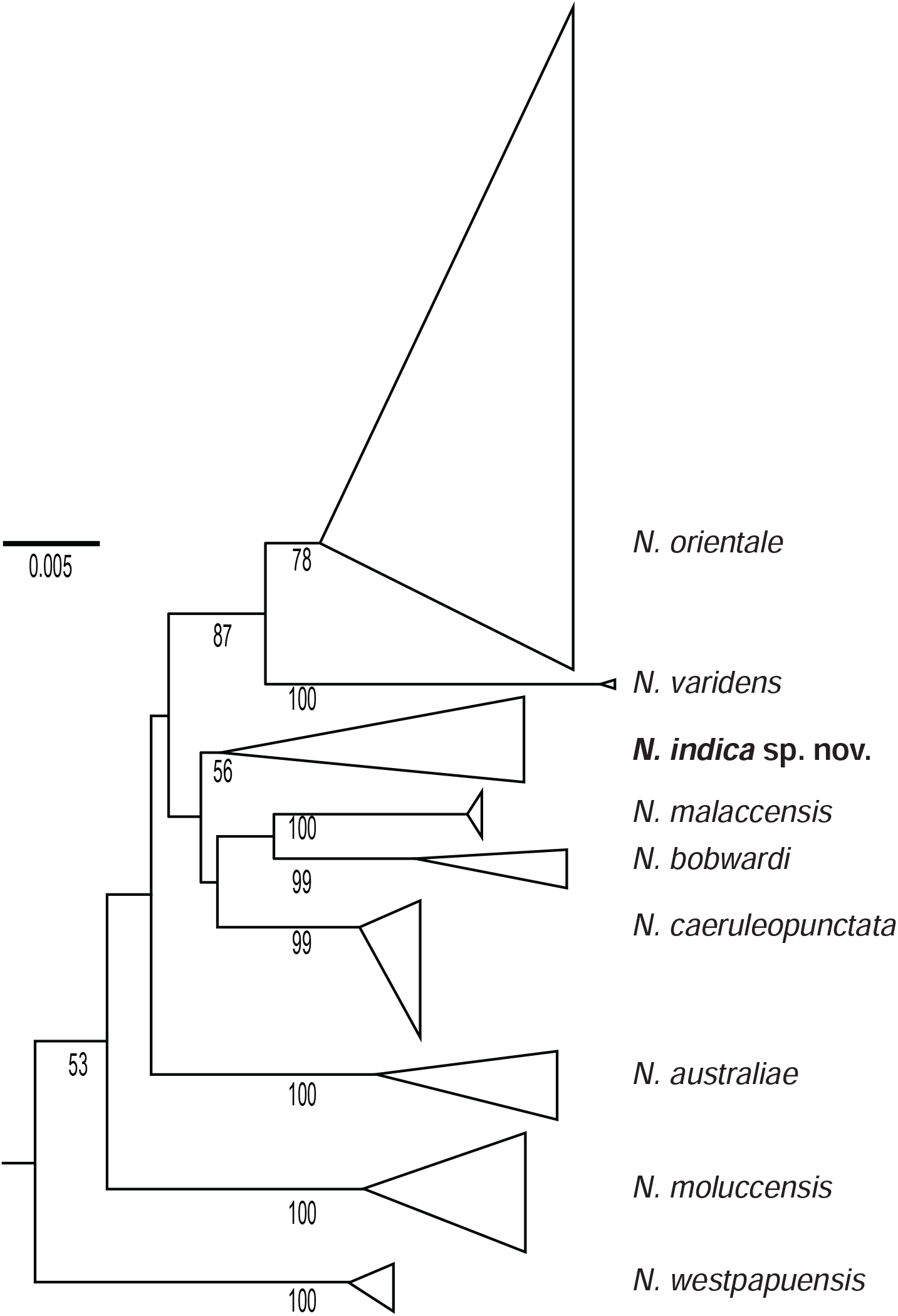
Blue-spotted maskray *Neotrygon* spp. Simplified maximum-likelihood tree of concatenated *CO1* + *cytochrome b* gene fragments, rooted by *N. trigonoides* ^15^ showing the placement of the *N. indica* sp. nov. lineage. Terminal branches were collapsed into triangles whose width is proportional to the number of individuals sequenced and whose depth represents the genetic variability within a species. Numbers at a node are bootstrap scores, from 600 bootstrap resampling runs under Mega6 ^27^.

Estimates of genetic distance between pairs of species in the Indian Ocean blue-spotted maskray group ranged from 0.020 to 0.025 at the *CO1* locus and from 0.020 to 0.033 at the *cytochrome b* locus (Table 3). These values were all well above the highest estimates of genetic distance within a species (respectively, 0.012 and 0.014; Table 3). Therefore, the four species albeit closely related remained distinct from one another using either the standard *CO1* barcode or the *cytochrome-b* marker.

**Table 3.** Blue-spotted maskray *Neotrygon* spp. from the Indian Ocean and geographically adjacent Andaman Sea. Estimates (mean ± *SD*) of nucleotide distances between pairs of, and within blue-spotted maskray species at loci *CO1* and *cytochrome b*.

| Marker,<br>Species | Within-species | Species |  |  |
| --- | --- | --- | --- | --- |
|  |  | *N. indica* | *N. bobwardi* | *N. caeruleopunctata* |
| *CO1* |  |  |  |  |
| *N. indica* | 0.012±0.003 | - |  |  |
| *N. bobwardi* | 0.009±0.003 | 0.025±0.006 | - |  |
| *N. caeruleopunctata* | 0.002±0.001 | 0.022±0.007 | 0.024±0.007 | - |
| *N. malaccensis* | 0.001±0.001 | 0.020±0.006 | 0.020±0.006 | 0.022±0.008 |
| *cytochrome b* |  |  |  |  |
| *N. indica* | 0.014±0.004 | - |  |  |
| *N. bobwardi* | 0.006±0.002 | 0.033±0.007 | - |  |
| *N. caeruleopunctata* | 0.002±0.001 | 0.020±0.005 | 0.026±0.006 | - |
| *N. malaccensis* | 0.000±0.000 | 0.026±0.006 | 0.026±0.009 | 0.021±0.007 |

### Mitochondrial DNA sequences as reliable taxonomic characters

Here, useful characters to distinguish *N. indica* sp. nov. from its blue-spotted maskray congeners derived from, in part (1) the analysis of spot patterns and, more accurately (2) molecular genetics. Although spot patterns proved somewhat helpful to tentatively characterize species including *N. indica* sp. nov., they were not fully diagnostic. Therefore, we chose to primarily base the description and diagnosis of the new species on the more powerful nucleotide sequence of its mitochondrial DNA, as done previously with other species of the genus *Neotrygon* ^7,2,15^. We have shown that meristics and morphometrics the way they were used in taxonomic descriptions of cryptic species in the blue-spotted maskray complex ^14^ hardly provided any diagnostic character ^7^. Part of the problem may stem from the fact that so far individual values of meristic and morphometric parameters have not been made available to the community except for a few type specimens ^14^.

## Taxonomy

*Neotrygon indica* sp. nov. (Fig. 1A, B; Fig. 2); urn:lsid:zoobank.org:act:99F4F1A4-D5F6-4379-BC4E-7FB1E74CFA58. *Trygon kuhlii* ^5^; presumably clade *Neotrygon kuhlii 3* of Naylor et al. ^18^; *Neotrygon kuhlii* ^15^; *Neotrygon kuhlii* haplogroup *I* ^10,11^; *Neotrygon kuhlii* clade *8* ^6^; Clade *I* ^9^; Indian-Ocean maskray ^7^. Also BIN number BOLD:AAA5611 in BOLD.

### Material examined

All specimens examined to characterize spot patterns are listed in Table 1. Specimens characterized morphometrically are listed in Table 4. The list of specimens characterized by their nucleotide sequence at mitochondrial loci *CO1* and *cytochrome b* is provided as appendix A of Borsa et al. ^19^. Additional *N. indica* sp. nov. specimens sequenced at the *CO1* and *cytochrome b* loci are listed in Table 2.

**Table 4.** Morphometric measurements ^14^ for four *Neotrygon indica* sp. nov. specimens from the eastern coast of India (holotype, paratype, NITT1, NITT2) and one specimen from Tanzania (IRD-20170720-A). All measurements, except disc width and number of caudal stings expressed as percentage of disc width.

| Parameter | Specimen |  |  |  |  |
| --- | --- | --- | --- | --- | --- |
|  | Holotype | Paratype | NITT1 | NITT2 | IRD-20170720-A |
| Disc width (mm) | 236 | 213 | 314 | 307 | 300 |
| Total length | 169.5 | 183.1 | 191.1 | 198.7 | - |
| Disc length | 80.5 | 84.5 | 79.6 | 81.4 | 80.4 |
| Distance from tip of snout to pectoral fin insertion | 69.9 | 70.4 | 73.9 | 76.5 | 71.9 |
| Distance from tip of snout to eye | 15.3 | 16.0 | 15.9 | 16.3 | 13.8 |
| Mouth width | 10.6 | 8.5 | 8.3 | 9.1 | 10.0 |
| Nostril length | 6.8 | 4.7 | 5.1 | 5.9 | 4.2 |
| Distance between nostrils | 11.9 | 14.1 | 9.6 | 10.7 | 7.5 |
| Head length | 40.3 | 39.9 | 39.8 | 42.3 | 40.8 |
| Eye length | 6.4 | 6.6 | 6.4 | 6.5 | 5.4 |
| Orbit diameter | 10.2 | 9.4 | 7.6 | 7.8 | 7.1 |
| Inter orbital width | 7.6 | 7.0 | 7.3 | 7.2 | 7.1 |
| Inter ocular width | 16.9 | 18.8 | 12.7 | 13.0 | 16.1 |
| Spiracle length | 7.6 | 8.0 | 7.6 | 7.2 | 7.1 |
| Interspiracular width | 14.8 | 15.0 | 12.7 | 14.3 | 15.6 |
| Orbit and spiracle length | 9.3 | 14.1 | 19.7 | 13.0 | 10.0 |
| Orbit to pectoral fin insertion | 55.9 | 46.9 | 51.0 | 53.7 | 52.2 |
| Nasal curtain length | 5.9 | 4.7 | 6.4 | 6.5 | 4.6 |
| Nasal curtain width | 9.3 | 8.5 | 8.9 | 9.1 | 9.4 |
| Width of 1<sup>st</sup> gill slit | 4.2 | 3.3 | 3.2 | 3.3 | 3.3 |
| Width of 3<sup>rd</sup> gill slit | 3.0 | 3.3 | 3.2 | 3.3 | 3.8 |
| Width of 5<sup>th</sup> gill slit | 1.7 | 2.3 | 2.2 | 2.3 | 2.1 |
| Distance between 1<sup>st</sup> and 5<sup>th</sup> gill slits (left) | 13.6 | 14.1 | 19.1 | 19.5 | 14.2 |
| Distance between 1<sup>st</sup> and 5<sup>th</sup> gill slits (right) | 13.6 | 14.1 | 13.4 | 13.0 | 14.2 |
| Distance between 1<sup>st</sup> gill slits | 16.9 | 16.9 | 14.9 | 16.2 | 17.5 |
| Distance between 5<sup>th</sup> gill slits | 11.0 | 9.5 | 8.3 | 8.4 | 9.6 |
| Pre-nasal length of snout | 9.3 | 5.7 | 9.6 | 9.8 | 5.6 |
| Distance from tip of snout to cloaca origin | 62.7 | 67.6 | 70.1 | 71.7 | 71.8 |
| Distance from pectoral fin insertion to base of 1<sup>st</sup> sting | 44.5 | 37.6 | 38.2 | 35.8 | - |
| Distance from origin of cloaca to base of 1<sup>st</sup> sting | 42.4 | 44.6 | 43.0 | 42.3 | - |
| Cloaca length | 7.2 | 6.6 | 7.3 | 7.5 | 4.6 |
| Tail width at axil of pelvic fin | 8.5 | 9.4 | 8.3 | 9.8 | 9.4 |
| Tail width at base of 1<sup>st</sup> sting | 2.3 | 6.6 | 3.8 | 4.6 | - |
| Tail height at axil of pelvic fin | 5.5 | 6.6 | 4.8 | 4.9 | 4.6 |
| Tail height at base of 1<sup>st</sup> sting | 2.6 | 2.3 | 2.5 | 2.9 | - |
| Pelvic fin (embedded) length | 14.4 | 13.6 | 15.9 | 14.7 | 14.7 |
| Width across pelvicfin base | 19.9 | 18.8 | 19.1 | 17.9 | 22.1 |
| Greatest width across pelvic fin | 33.9 | 42.3 | 41.4 | 44.3 | 43.0 |
| Clasper length from pelvic axil | - | - | 15.6 | 16.6 | - |
| Post-cloacal clasper length | - | - | 20.4 | 22.8 | - |
| Tail width at base of sting | 3.4 | 3.8 | 7.6 | 7.8 | - |
| Tail height at base of sting | 2.5 | 2.8 | 3.8 | 3.9 | - |
| Disc thickness | 14.8 | 14.1 | 14.3 | 13.7 | 13.5 |
| Number of caudal stings | 1 | 1 | 2 | 2 | - |
| Length of 1<sup>st</sup> sting | - | 14.1 | 16.2 | 15.0 | - |
| Length of 2<sup>nd</sup> sting | - | - | 14.3 | 18.6 | - |

### Types

A female specimen 236 mm disc width (Fig. 1A) from the Gulf of Mannar, Tamil Nadu (9.12°N 79.46°E), collected at the Inico Nagar, Tuticorin fish landing centre by RK and Satish on 13 January 2017 is here designated as the holotype of *Neotrygon indica* sp. nov. The specimen was deposited by APK at the MBRC in Chennai, India on 12 May 2017. The specimen has registration no. ZSI/MBRC/F.1495

A female specimen 213 mm disc width (Fig. 2) fished from the Gulf of Mannar, Tamil Nadu (9.12°N 79.46°E) using bottom-set gillnets, collected by RK on 25 January 2017 is here designated as the single paratype of the new species. This specimen, labelled CIFEFGB/NKGM1 was subsequently deposited by APK at the Fish Genetics and Biotechnology laboratory, ICAR-Central Institute of Fisheries Education, Mumbai, India.

### Description of holotype

Ocellated blue-spot, dark-spot and dark-speckle counts on the dorsal side of the holotype are presented in Table 1. The meristic details of the holotype, which were scored from X-ray photographs, were the following: number of pectoral-fin radials 106-108 including propterygion 47-51, mesopterygion 12 and metapterygion 45-47. Morphometric measurements on the holotype are presented in Table 4.

### Measurements on paratype

Similarly, spot pattern details on the dorsal side of the paratype are presented in Table 1 and its meristic details were scored from X-ray photographs. The number of pectoral-fin radials was 108-110 including propterygion 45-47, mesopterygion 14-15 and metapterygion 47-50. Morphometric measurements are presented in Table 4.

### Diagnosis

*Neotrygon indica* sp. nov. from India is differentiated from all species of the blue-spotted maskray described so far (*N. australiae*, *N. bobwardi, N. caeruleopunctata*, *N. malaccensis*, *N. moluccensis*, *N. orientale*, *N. vali*, *N. varidens* and *N. westpapuensis*) by a combination of low number of medium-sized ocellated blue spots, total absence of large ocellated blue spots, high number of dark speckles, frequent occurrence of a few dark spots, and conspicuous occipital mark. However, this diagnosis did not apply to the two specimens sampled from Zanzibar whose spot patterns were examined. *Neotrygon indica* sp. nov. is distinct from *N. kuhlii* and *N. trigonoides* by the absence of the pair of conspicuous scapular blotches characteristic of these two species ^15,16^, although specimens of *N. indica* sp. nov. often possess a pair of diffuse darker marks in the scapular region.

Fifteen out of 19 individuals of *N. indica* sp. nov. sequenced at the *CO1* locus possessed T at nucleotide site 607 of the *CO1* gene, a character that was otherwise present in only two out of 130 *N. ońentale* individuals and absent in all other cryptic species of the blue-spotted maskray. *Neotrygon indica* sp. nov. was further distinguished from other species previously under *N. kuhlii* by A at nucleotide site 70 of the *cytochrome b* gene (based on 5 individuals from Tanzania). Eight out of 11 individuals of *N. indica* sp. nov. also possessed T at nucleotide site 284 of the *cytochrome b* gene, a character that was otherwise present in only one out of 77 *N. ońentale* individuals and absent in all other cryptic species of the blue-spotted maskray.

### Distribution

The type locality of *N. indica* sp. nov. is the Gulf of Mannar in Tamil Nadu on the eastern coast of the Indian sub-continent. Based on the collection of voucher specimens from present study, the distribution of *N. indica* sp. nov. includes the coasts of Andhra Pradesh and Tamil Nadu states of India, from approximately 17.7°N to 8.8°N. Based on the material genetically examined thus far, the distribution of *N. indica* sp. nov. also includes the Indian coast of the Laccadives Sea (Kerala) and Tanzania.

### Etymology

Epithet *indica* is the latin feminine adjectival form of the name of the country of type locality, India.

### Proposed vernacular names

Indian-Ocean blue-spotted maskray (English); Neeli Nishan Pakat (Hindi); Pulli Thirukhai (Tamil); Raie pastenague masquée à points bleus de l’océan Indien (French).

